# Characterization of the First Turtle Organoids: A Model for Investigating Unique Adaptations with Biomedical Potential

**DOI:** 10.1101/2023.02.20.527070

**Authors:** Christopher Zdyrski, Vojtech Gabriel, Thea B. Gessler, Abigail Ralston, Itzel Sifuentes-Romero, Debosmita Kundu, Sydney Honold, Hannah Wickham, Nicholas E. Topping, Dipak Kumar Sahoo, Basanta Bista, Jeffrey Tamplin, Oscar Ospina, Pablo Piñeyro, David K. Meyerholz, Karin Allenspach, Jonathan P. Mochel, Nicole Valenzuela

## Abstract

Painted turtles are remarkable for their well-developed freeze tolerance and associated resilience to hypoxia/anoxia, oxidative stress, and ability to supercool. They are, therefore, an ideal model for biomedical research on hypoxia-induced injuries (including strokes), tissue cooling during extensive surgeries, and organ cryopreservation. Yet, the seasonal reproduction and slow maturation of turtles hinder basic and applied biomedical research. To overcome these limitations, we developed the first adult stem cell-derived turtle hepatic organoids, which provide 3D self-assembled structures that mimic their original tissue and allow for *in vitro* testing and experimentation without constantly harvesting donor tissue and screening offspring. Our pioneering work with turtles represents the first for this vertebrate Order and complements the only other organoid lines from non-avian reptiles, derived from snake venom glands. Here we report the isolation and characterization of hepatic organoids derived from painted, snapping, and spiny softshell turtles spanning ∼175 million years of evolution, with a subset being preserved in a biobank. Morphological and transcriptomics revealed organoid cells resembling cholangiocytes, which was then compared to the tissue of origin. Deriving turtle organoids from multiple species and life stages demonstrates that our techniques are broadly applicable to chelonians, permitting the development of functional genomic tools currently missing in most herpetological research. When combined with genetic editing, this platform will further support studies of genome-to-phenome mapping, gene function, genome architecture, and adaptive responses to climate change, among others. We discuss the unique abilities of turtles, including their overwintering potential, which has implications for ecological, evolutionary, and biomedical research.

**SIGNIFICANCE:** Here we developed the first turtle-derived organoid biobank from the liver of multiple chelonians with a subset characterized via histology, RNA sequencing transcriptomics, single-nuclei RNA sequencing, and transmission electron microscopy. Furthermore, we discuss the potential of the 3D organoid model to investigate unique physiological adaptations of turtles which could unravel the molecular mechanisms underlying their overwintering capacity, opening the door for *in vitro* biomedical studies relevant to hepatic ischemia-reperfusion injury to organ cryopreservation, beyond fundamental ecology and evolution. This organoid biobank represents a novel resource for the scientific community to support research regarding the unique adaptations of this understudied Order of vertebrates.

## INTRODUCTION

Turtles are an ancient and enigmatic group of reptiles recognized for their distinctive morphology, longevity, tolerance to anoxia, and diverse sex-determining systems. Importantly, the study of turtles is both time-sensitive and critical, due to their at-risk status due to anthropogenic global environmental change and habitat loss, among others (Lovich et al., 2018; Rhodin et al., 2018). Because of their biology and phylogenic position, turtles hold clues to unlock numerous biological mysteries, including current biomedical questions. Indeed, turtles are an emerging model for ecology, evolution, and human health (Valenzuela, 2009), currently studied to understand physiology, life histories, chromosome evolution, as ecotoxicology sentinels, and to decipher biological pathways for sexual development and reproduction (Bista & Valenzuela, 2020; Chaousis et al., 2021; Congdon et al., 2022; Lee et al., 2020; Mizoguchi et al., 2022; Montiel et al., 2016; Sabath et al., 2016; Thépot, 2021). But while reptile genomics is thriving, reptilian transgenics remains challenging despite pioneering *in vivo* gene editing in anole lizards (Rasys et al., 2019) and recent work in mourning geckos (Lozito et al., 2021). One reason is that turtle research is currently hindered due to the scarcity of *in vitro* tools, and additionally impeded by the slow maturation and seasonal reproduction of chelonians. Thus, developing methods that overcome these bottlenecks and increase their use in basic and applied research to study their remarkable adaptations is overdue. Stem cell-derived organoids (Sato et al., 2009) are an attractive model for functional genomics as they form complex 3D structures that recapitulate the microanatomy and physiology of their tissue of origin (de Souza, 2018; Lehmann et al., 2019). Unlike conventional 2D cell cultures, adult stem cell-derived organoids have several advantages, including being composed of multiple epithelial cell types also found in the tissue of origin, being able to continuously be expanded, and being able to self-renew and self-organize (Fatehullah et al., 2016).

Organoid technology has expanded recently from commonly used mice and human models to canines (Ambrosini et al., 2020; Chandra et al., 2019; Gabriel et al., 2022; Minkler et al., 2021; Mochel et al., 2018; Nantasanti et al., 2015) and few other vertebrates. However, the only reptilian (sensu lato) organoids (i.e., excluding those derived from chicken intestines (Zhao et al., 2022)), were generated from snake venom glands (Post et al., 2020), representing only a fraction of the vast Tree of Life. Expanding the taxonomic coverage of the 3D organoid technology will leverage unique adaptations that evolved in reptiles and other non-model species, broadening potential applications of this technology. Here we report the generation of hepatic organoids from painted turtles (*Chrysemys picta*), snapping turtles (*Chelydra serpentina*), and spiny softshell turtles (*Apalone spinifera*), and their characterization via histological staining, RNA-seq transcriptomics, single-nuclei RNA-seq, and transmission electron microscopy.

Furthermore, painted turtles are well-adapted to overwintering conditions by supercooling their body and surviving in the ensuing hypoxic and ischemic conditions (Storey & Storey, 2017). They are one of the most anoxia-tolerant tetrapods (Bradley Shaffer et al., 2013), which, along with the slider turtle (*Trachemys scripta*), can survive for weeks without oxygen (Krivoruchko & Storey, 2015). Because the liver is critical to the adaptive defense underlying their supercooling capacity and tolerance to anoxia (Dinkelacker et al., 2005; Storey, 2006), the development of hepatic organoids opens the door for new biomedical research.

Indeed, certain human diseases cause hypoxia in vital organs, as reported in liver cirrhosis, an end-stage disease caused by chronic injuries to the hepatic tissue by drugs, alcohol, infections, and genetic disorders (Roth & Copple, 2015). Acute injury to the liver can consequentially cause the upregulation of hypoxia-inducible factors (HIFs) to maintain homeostasis (Moon et al., 2009; Roth & Copple, 2015). Furthermore, hepatic ischemia-reperfusion injury (IRI), a major clinical complication during liver transplantation, severe trauma, vascular surgery, and hemorrhagic shock (Kaltenmeier et al., 2022), seems improved by cold machine-perfusion of organs before transplantation (Jia et al., 2020). Thus, understanding the capability of turtle hepatic organoids to survive hypoxia and anoxia may aid in discovering important proteins possibly implicated which would aid the future development of treatment options for IRI. For instance, antifreeze glycoproteins prolonged the survival of mouse intestinal organoids when incubated at 4°C for up to 72 hours (Huelsz-Prince et al., 2019). Turtle hepatic organoids should help illuminate their unique adaptative strategies to overwintering which could benefit human organ preservation medicine. Overall, the development of organoids from non-model species, such as turtles, can greatly impact biomedical research by exploiting a myriad of adaptations found across the Tree of Life.

## RESULTS

### Growth and expansion of hepatic organoids

Here we report the culture of the first turtle-derived organoids from the liver of juvenile spiny softshell turtle (*Apalone spinifera*) and snapping turtle (*Chelydra serpentina*), as well as embryonic, hatchling, and adult painted turtles (*Chrysemys picta*) (**Fig. 1*B* and *C***). Our optimization of the isolation procedure for turtle hepatic organoids eliminated unnecessary steps typically done when attempting to isolate intestinal stem cells (Gabriel et al., 2022), which expedited the final experimental protocol (**Fig. 1*A***) and included media supplementation with prostaglandin E2 (PGE2) 9.93 nM. A biobank was generated that contains hepatic organoid lines from *C. serpentina* (*n*=1), and *C. picta* (*n*=1 embryonic, *n*=3 hatchling, and *n*=3 adult) (**Table S1**). Organoid lines withstood multiple passages, with the embryonic organoids of painted turtles being passaged a total of 14 times (**Table S1**). Importantly, six organoid lines from the biobank were successfully thawed, re-cultured, and expanded, including juvenile *C. serpentina* (*n*=1), plus embryonic (*n*=1), hatchling (*n*=1), and adult (*n*=3) *C. picta* (**Table S1**). Although *A*. *spinifera* organoids were successfully isolated, they typically did not proliferate into larger numbers after the second or third passage as needed, such that samples were collected solely for histological characterization, while further optimization of media and temperature is warranted to improve their long-term culture in the future.

**Fig. 1:**
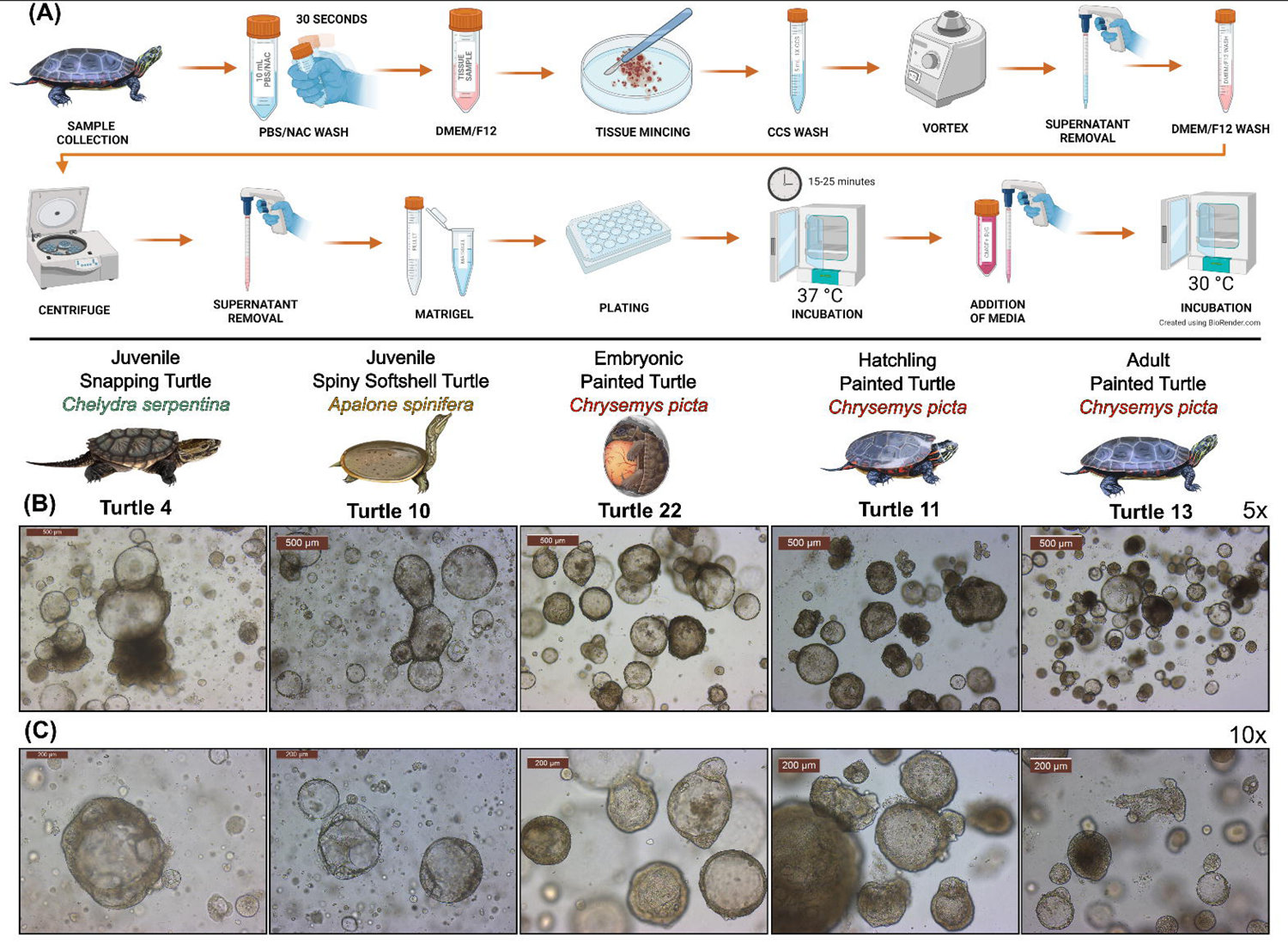
Morphological characterization and isolation optimization for turtle hepatic organoids. Culture protocol and characterization of turtle hepatic organoids. (A) Minimal workflow to isolate and culture turtle organoids, from tissue collection, mincing, washing, plating, and incubating the turtle liver tissue. (B-C) Light microscopy images of organoids derived from the liver of a spiny softshell turtle (*Apalone spinifera*), the snapping turtle (*Chelydra serpentina*), as well as embryonic, hatchling, and adult painted turtles (*Chrysemys picta*) at 5X (B) and 10X (C) magnification.

### Morphological organoid characterization

The morphology of all the turtle hepatic organoids under bright field microscopy mostly resembled cystic spheroids, although darker spheroids with a thicker organoid border were also observed (**Fig. 1*B* and *C***). Additionally, *A. spinifera* hepatic organoids typically displayed a thinner organoid border and visible vacuoles in the organoid compared with *C. serpentina* and *C. picta* (**Fig. 1*C***). Hematoxylin and eosin (H&E) staining identified the structure of the cells and microanatomy present (**Fig. 2*A*-*D***), as well as identifying hepatocytes (**Fig. 2*D***) in the tissues of origin. Staining with Alcian Blue uncovered mucin production within *C. picta* hatchling hepatic organoids on the apical side of the lumen (**Fig. 2*G* and *H***), revealing hepatic secretory cells which we hypothesize to be cholangiocytes. Periodic Acid-Schiff (PAS) was also used to identify glycogen in the tissue samples (**Fig. 2*E***). Analysis with transmission electron microscopy (TEM) identified cellular components including the nucleus, nucleolus, and mitochondria in the liver tissues (**Fig. 2*F***) and hepatic organoids (**Fig. 2*K***). Additionally, in the adult hepatic organoids, TEM identified the secretion of mucin (**Fig. 2*I* and *J***), consistent with the Alcian Blue staining of hatchling hepatic organoids.

**Fig. 2:**
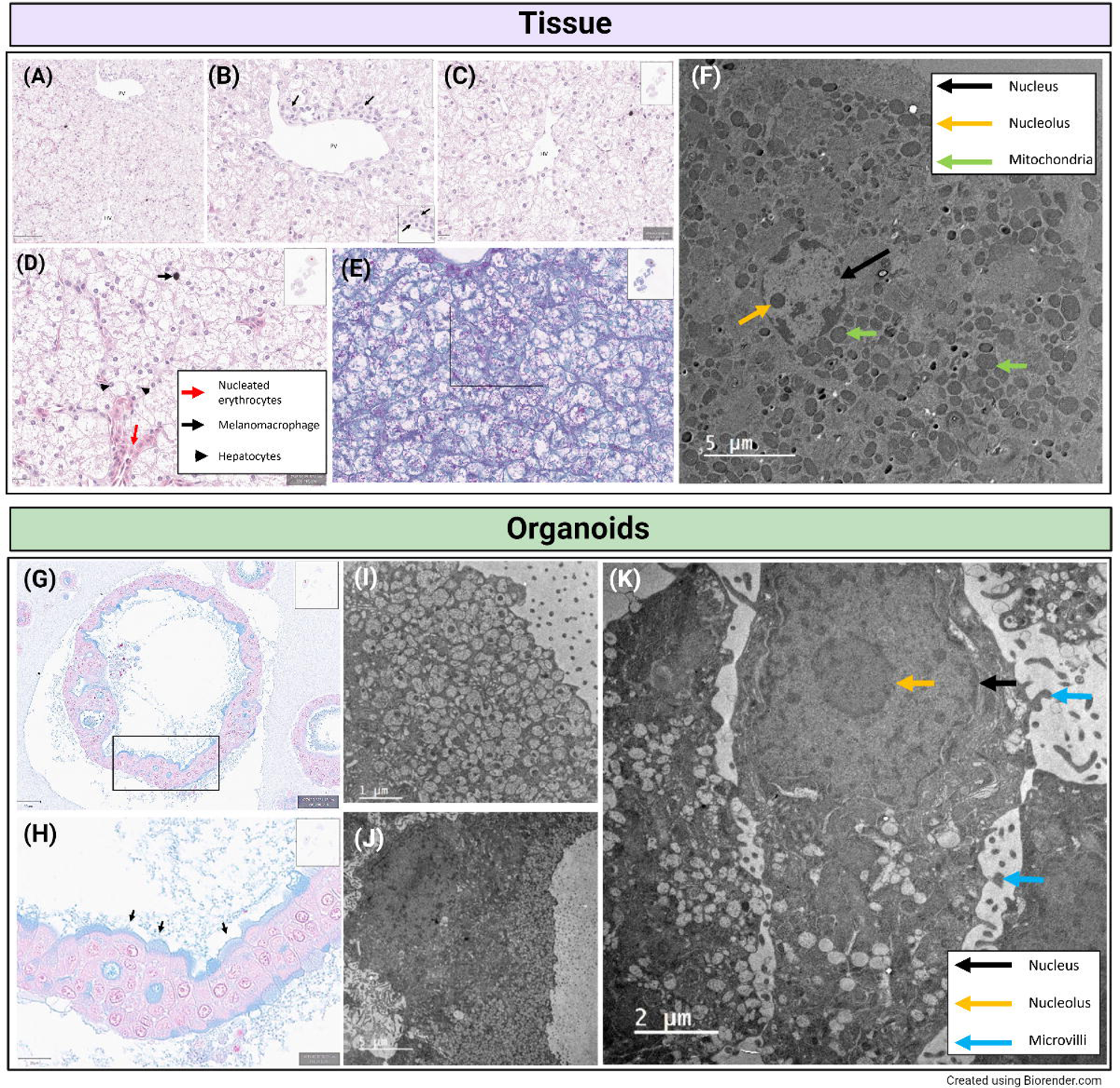
Histopathology staining and transmission electron microscopy of painted turtle hepatic tissues and organoids. Overview of turtle liver histology from a hatchling painted turtle (*Chrysemys picta*). (A-C) Key histologic landmarks identified using hematoxylin and eosin (H&E) in the turtle liver tissue include the portal venule (Pv) and terminal hepatic venule (Hv). In the portal region, a small bile duct is seen (B, inset and arrows). (D) Nucleated erythrocytes are seen within sinusoids (red arrows) and melanomacrophages (black arrow) are readily detected by their cytoplasmic pigment (black arrows). Hepatocytes (black arrowheads) often have round nuclei that are eccentrically and basolaterally located in the cytoplasm near the sinusoidal border. Hepatocytes are distended and have increased rarefaction of the cytoplasm, which parallels the presence of PAS+ glycogen (E, magenta coloration). (F) Transmission electron microscopy (TEM) of turtle liver tissue with arrows identifying structures of interest (nucleus = black, nucleolus = yellow, green = mitochondria). (G, H) Alcian blue histochemical stain of hatchling *Chrysemys picta* hepatic organoids showed AB+ mucins along the apical border of cells lining the lumina (inset and arrows). Note also that the cells have round nucleus that has eccentric, basolateral localization in the cytoplasm. (I, J) TEM images of turtle hepatic organoids show mucin granules in apical cytoplasm, confirming AB + staining. (K) TEM of the turtle hepatic organoids with arrows identifying structures (nucleus = black, nucleolus = yellow, blue = microvilli). Scale bars are in µm.

### Bulk stranded RNA-seq analysis of hatchling and adult painted turtles for hepatic organoids and liver tissue

For *C. picta*, over 94% of RNA-seq reads from each library (tissue and organoids from both hatchlings and adults) mapped as pairs to the reference genome (Chrysemys_picta_BioNano-3.0.4). For *C. serpentina*, the mapping rate to the painted turtle reference genome was lower (88% of pairs for the organoid and 84% of pairs for the tissue), likely due to divergence of their genomes during ∼105My since they evolutionarily split from each other, which impairs cross-species mapping (**Table S2**). We used *C. picta* genome as a reference because the *C. serpentina* genome assembly (Das et al., 2020), is not fully annotated and is much more fragmentary. A principal components analysis (PCA) of the normalized gene model counts for liver organoid and liver tissue transcriptomes in painted turtles indicated strong clustering by both age and sample type (**Fig. 3*A***), as well as interspecific differences when snapping turtle data were included (**Fig. 3*B***). In *C. picta* alone (**Fig. 3*A***), PC1 primarily captured variation (47.10%) due to sample type (tissue vs. organoid), while PC2 primarily captured variation (9.32%) due to age (hatchling vs. adult). When including *C. serpentina* (**Fig. 3*B***), PC1 still captured variation (34.76%) due mostly to sample type (tissue vs. organoid), a relationship that was retained across species, while PC2 primarily captured variation (17.49%) due to species, and to a lesser degree by age (hatchling vs. adult) when examining *C. picta* clusters. The transcriptome assembly for painted turtles generated 54,050 gene models, of which 27,876 had a baseMean of > 50. Most genes (>26K) were expressed in both hepatic organoids and tissues, while 4.8K genes were uniquely expressed in hepatic organoids and 9K in liver tissue when hatchlings and adults were combined (**Fig. 3*C***). Similarly, when data were separated by life stage, the majority of genes (∼23K) were expressed in both organoids and tissues of hatchlings and adults, whereas only ∼1K genes were uniquely expressed in organoids of any age, and ∼1K or ∼2.6K genes were uniquely expressed in hatchling or adult tissue, respectively (**Fig. 3*D***). Although most genes were expressed in common, DEGs were detected between organoids and tissues (15,671 in adults and 14,711 in hatchlings). **Fig. 3*E*-*H*** illustrates the top 50 most divergently expressed genes between tissues and organoids (either up-[red] or downregulated [blue]). As expected across life stages, differentially expressed genes (DEGs) were detected between adult and hatchling organoids (884 DEGs), and between adult and hatchling tissues (3,839 DEGs).

**Fig. 3:**
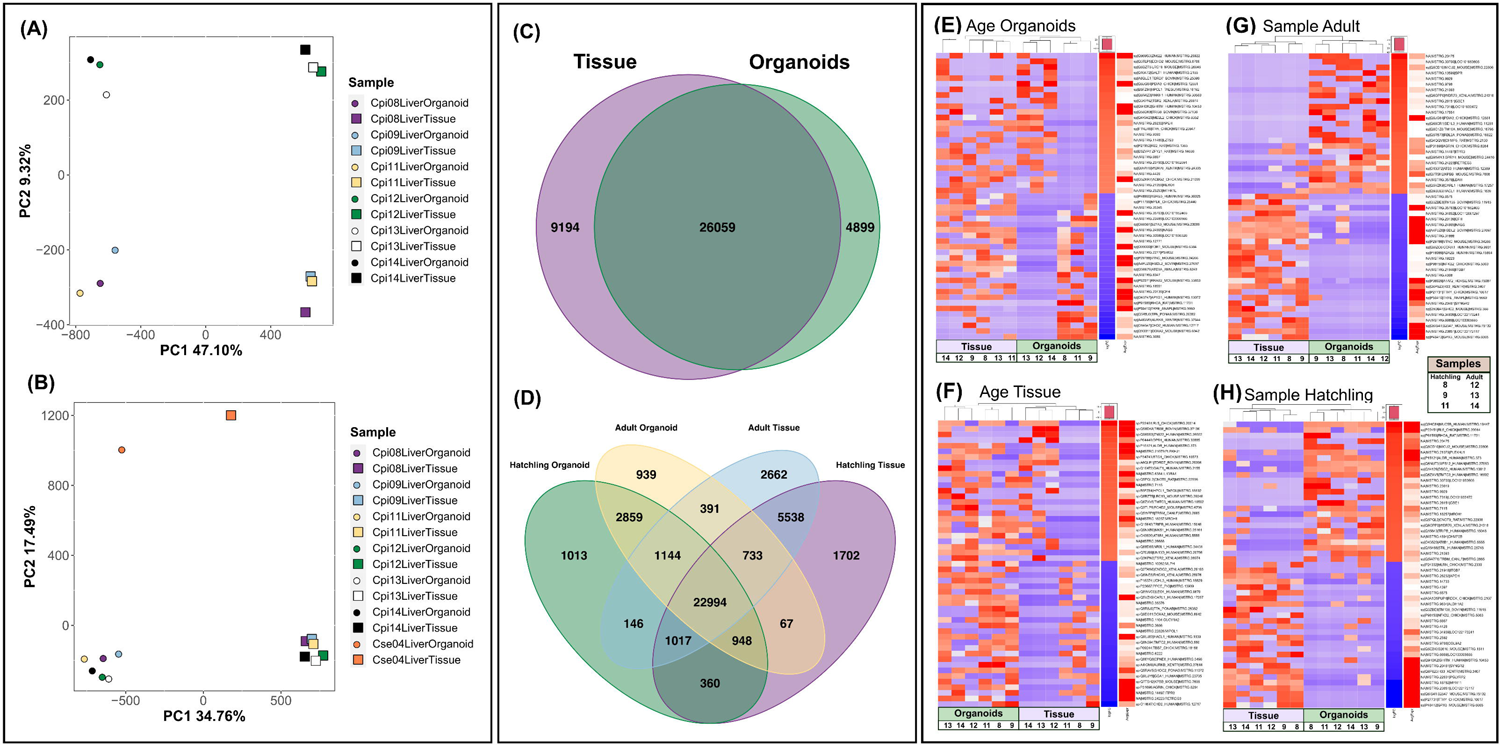
Transcriptomic characterization of turtle hepatic organoids and liver tissue of origin. Transcriptomic expression patterns of liver tissue and hepatic organoids. (A) A principal components analysis (PCA) displaying clustering of sample types for *Chrysemys picta* (circles = organoids; squares = tissues) (*C. picta* hatchlings *=* purple, blue, yellow; *C. picta* adults *=* green, white, black). (B) Inclusion of *Chelydra serpentina* (orange) in the PCA plot reveals species-specific differences. (C) Venn diagram identifying the number of shared and uniquely expressed genes in *C. picta* tissues (purple) and organoids (green) from hatchlings and adults combined. (D) Venn diagram identifying shared and uniquely expressed genes across *C. picta* sample types and life stages (hatchling organoid = green, adult organoid = yellow, hatchling tissue = purple, adult tissue = blue). Heatmaps of differentially-expressed genes (DEGs) in *C. picta* between age groups (hatchling vs. adult) and sample type (organoid vs. tissue). Specifically, heatmaps identified DEGs when comparing (E) age within organoids, (F) age within tissue, (G) sample type in adults, and (H) sample type in hatchlings.

### Annotation of C. picta expression differences across life stages

When searching the transcriptomes for genes involved in hepatic function, two important genes were upregulated in *C. picta* hatchling hepatic organoids compared with adult hepatic organoids (**Fig. 3*E***). The first is tissue-type plasminogen activator (*Tpa*) which is involved in plasminogen conversion into plasmin (a main enzyme responsible for clot breakdown) that is currently used in medical applications and was the only previously approved pharmacological treatment for restoring blood flow after a stroke occurred (Henderson et al., 2018). The second gene is transforming protein RhoA (*Rhoa*), which is involved in a pathway that promotes actin polymerization in the cell cytoskeleton (among other changes) and is thought to play a role in the anoxic overwintering ability of turtles, affecting the actin dynamics in the cell (Myrka & Buck, 2021). In contrast, genes that were upregulated in adult compared with hatchling hepatic organoids included protein disulfide-isomerase A3 (*Pdia3*) and enoyl-CoA hydratase domain-containing protein 2 (*Echd2*), both of which encode proteins secreted from primary human hepatocytes (Franko et al., 2019). Additionally, dual oxidase maturation factor 2 (*Doxa2*), was upregulated in hatchling compared with adult hepatic organoids, which is a gene found to be upregulated in the liver of the Chinese softshell turtle (*Pelodiscus sinensis*) under anoxic conditions, possibly due to increased reactive oxygen species (ROS) or disturbed mitochondrial function (Zhang et al., 2018).

In liver tissues (**Fig. 3*F***), upregulated genes in adult *C. picta* compared with hatchlings included 60S ribosomal protein L5 (*Rl5*), which is associated with the cellular mechanism that is responsible for translating mRNA to proteins (Hassan et al., 2021). Additionally, protein BTG1 (*Btg1*) was upregulated in all three adults compared with hatchling tissues, a gene encoding a protein with anti-proliferative function that prevents cell growth and lowers energy demands, which is upregulated in painted turtles under anoxic conditions (Keenan et al., 2015).

### Differentially expressed genes of interest between C. picta hepatic organoids and liver tissues from hatchlings and adults

In adult hepatic organoids, many DEGs were involved in glycogen production or storage (**Fig. 3*G***), two important liver functions that regulate energy metabolism. Upregulated genes in adult hepatic organoids compared with adult liver tissue included 1,4-alpha-glucan branching enzyme 1 (*Gbe1*), a glycogen branching enzyme, protein disulfide-isomerase A3 (*Pdia3*), which is secreted by hepatocytes (Franko et al., 2019) and responds to endoplasmic reticulum stress, helping modulate the folding of newly synthesized glycoproteins (Kondo et al., 2019). Compared with adult liver tissues, adult hepatic organoids also showed upregulation of two additional interesting genes. The first is tRNA methyltransferase 10 homolog A (*Tm10a*), a tRNA modification enzyme, that may be involved in the protective stress response (Wei & Tomizawa, 2018). The second is phosphorylase b kinase regulatory subunit beta (*Phkb*; *Kpbb*), which stimulates glycogen breakdown (S. Yang et al., 2018), and predicts poor prognosis in human hepatocellular carcinoma patients when downregulated, whereas it inhibits cell proliferation and induces apoptosis of tumor cells when artificially upregulated (W. Yang et al., 2019).

Finally, several genes were upregulated in hatchling hepatic organoids compared with hatchling tissues (**Fig. 3*H***). These include Mucin-5B (*Muc5b*), a mucin expressed in mammalian cholangiocytes (MacParland et al., 2018), transforming protein RhoA (*Rhoa*), which is involved in anoxic overwintering in the painted turtle (Myrka & Buck, 2021), and 1,4-alpha-glucan branching enzyme 1 (*Gbe1*), a glycogen branching enzyme. Several other DEGs were upregulated in hatchling hepatic organoids, including protein O-mannosyl-transferase TMTC3 (*Tmtc3*) which, when downregulated, reduces transcripts that are involved in degrading proteins (Racapé et al., 2011), and CCR4-NOT transcription complex subunit 9 (*Cnot9*), a member of the CCR4-NOT complex which helps maintain liver homeostasis via mRNA deadenylation in order to modulate the liver transcriptome (Takahashi et al., 2020).

### Enrichment analysis

The graphical results from treemap (**Supplemental Fig. 1-3**) illustrate the high degree of similarity in enrichment patterns between all groups in *C. picta* (hatchling tissue, hatchling organoid, adult tissue, adult organoid). The group terms obtained by treemap provide a supercluster representative, describing the general patterns of multiple related clusters of genes. While a deeper examination reveals more granular details of specific terms present, these supercluster terms allow for assisted identification of trends.

*For Biological Process GO terms* (**Supplemental Fig. 1**), hatchling and adult organoid groups shared the same top three supercluster terms: organonitrogen compound catabolic process, organic substance metabolic process, and nitrogen compound transport. Further, hatchling and adult tissues also shared the organic substance metabolic process with both organoid groups, and adult tissues also shared enrichment of organonitrogen compound catabolic process with hatchling and adult organoids. mRNA metabolic process was a top supercluster term for both hatchling and adult tissues. A third top supercluster term for hatchling tissues was cellular catabolic process.

*For Cellular Component GO terms* (**Supplemental Fig. 2**), ribonucleoprotein complex was a top supercluster term for all four groups. Adult and hatchling organoids shared the term organelle subcompartment, while hatchling tissues included the term intracellular organelle lumen, and adult tissues included the term organelle membrane and cytosol. The second-tier superclusters for all four groups overlapped in most of their terms including: endomembrane system, intracellular anatomical structure, membrane-enclosed lumen, organelle, and protein-containing complex. Both organoid groups also shared the supercluster term envelope.

*For Molecular Function GO terms* (**Supplemental Fig. 3**), among top supercluster terms shared by all four groups were mRNA binding, catalytic activity, structural constituent of ribosome, and structural molecule activity. Hatchling and adult organoids also shared the supercluster term hydrolase activity. An additional prominent supercluster term for hatchling organoids was catalytic activity, acting on a nucleic acid.

These biological processes, cellular components, and molecular functions, demonstrate the overall shared expression patterns in the top superclusters are seen between groups when comparing *C*. *picta* across age (hatchlings and adults) and sample type (organoids and tissues), highlighting the potential of the turtle hepatic organoid model as a useful *in vitro* tool for a variety of research topics.

### Single-nuclei RNA-seq reveals cell clusters in embryonic hepatic organoids

After transcriptomes of hatchling and adult *C. picta* hepatic organoids had been characterized, we attempted and were successful in growing embryonic turtle hepatic organoids, whose transcriptome was characterized using single-nuclei RNA-seq (snRNA-seq) to identify relevant cell clusters. This dataset is composed of 387,570,615 sequenced reads derived from 21,384 single nuclei samples with a total of 18,852 features. Overall, 95.3% of the reads mapped to the painted turtle reference genome. A total of 8 distinct cell clusters were identified in the embryonic *C. picta* hepatic organoids (**Fig. 4*A***) with one gene from each cluster used to visualize expression across all cells (**Fig. 4*B***). Expression of each cluster was compared to known liver cell types (**Fig. 4*C***) and specific cholangiocyte cell types (**Fig. 4*D***) were used to further characterize the expression seen across clusters. Based on these known cell markers, all clusters had a strong cholangiocyte signature (KRT8 and KRT18) while a subset of cells also had limited expression of markers found in progenitor-associated cells (ALCAM and WWTR1). Upon further division of specific cholangiocyte cell types, markers for mature cholangiocytes were the most prominent for all clusters. Finally, a heatmap displayed the expression of the top genes (upregulated = yellow, downregulated = purple) in each cluster (**Fig. 4*E***).

**Fig. 4:**
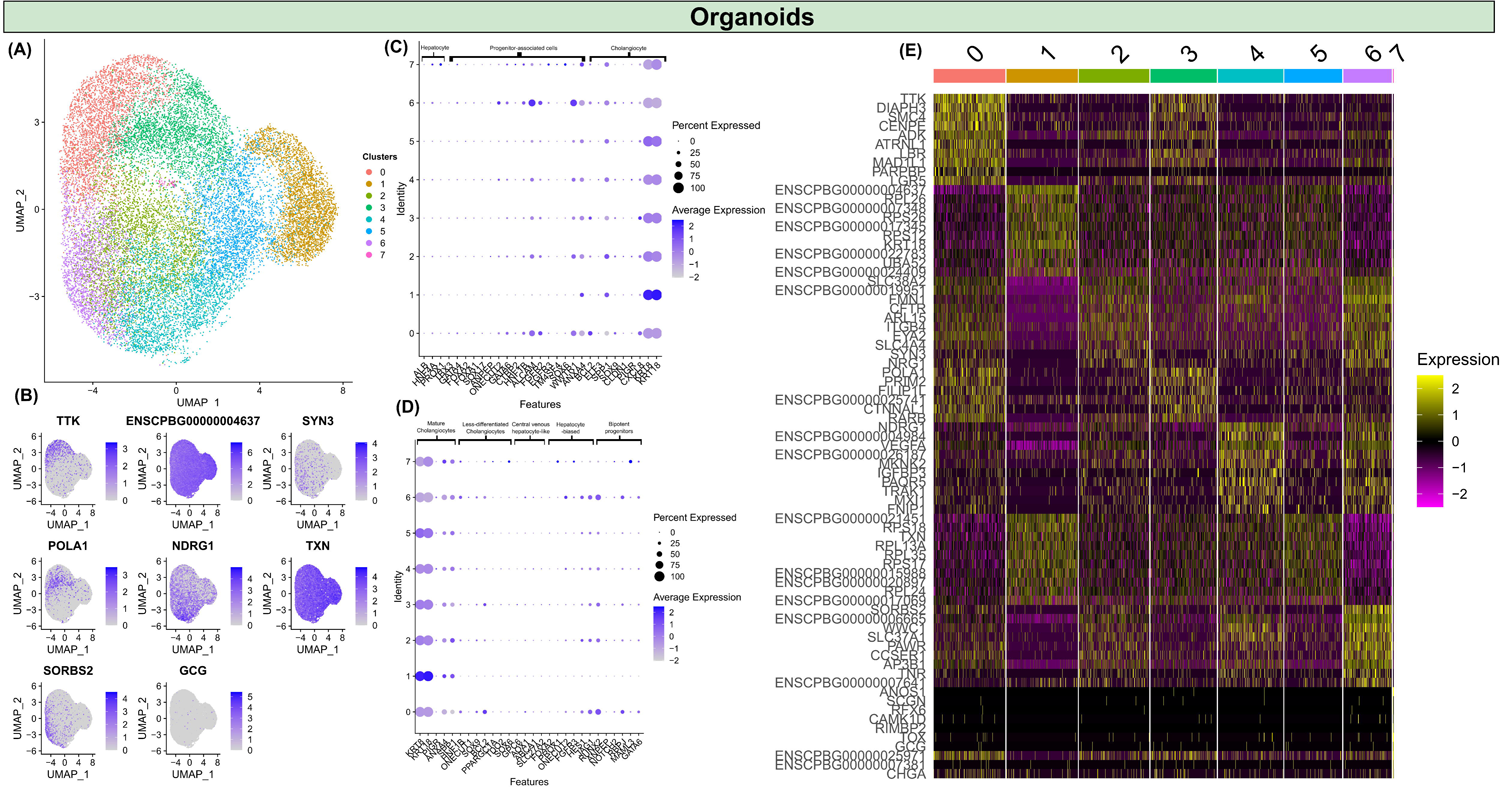
Single-nuclei RNA-seq analysis of embryonic *Chrysemys picta* hepatic organoids. Identification of cell clusters using snRNA-seq. (A) Unannotated UMAP showing the 8 distinct cell clusters identified in embryonic *Chrysemys picta* hepatic organoids. (B) One gene expressed in each cluster displaying the expression intensity across clusters. (C) Expression of known genes representing hepatocytes, progenitor-associated cells, and cholangiocytes for each cluster (T. S. Andrews et al., 2021). (D) Expression of known genes representing mature cholangiocytes, less-differentiated cholangiocytes, central venous hepatocyte-like, hepatocyte-biased, and bipotent progenitor cells for each cluster (T. S. Andrews et al., 2021). (E) Heatmaps identifying up to the top 20 markers for each cluster (upregulated = yellow and downregulated = purple, with respect to each other).

## DISCUSSION

### The promise of 3D turtle organoid technology

Scientific interest in reptilian genomics has expanded greatly since the sequencing of the green anole *Anolis carolinensis* (Alföldi et al., 2011), and the taxonomic scope of studies has expanded greatly from single representatives of major non-avian reptilian lineages (Janes et al., 2008) due to mounting genomic resources. As expected, reptiles have lagged behind traditional biomedical research models such as *Caenorhabditis elegans*, *Drosophila melanogaster*, *Danio rerio,* and *Mus musculus* for a long time (Kim et al., 2020). While there are good reasons for this, including the small size, fast life cycle, and high genetic or physiological similarities to humans of some traditional models (Cornet et al., 2018), the unique evolutionary adaptations of non-model taxa provide an opportunity to answer fundamental questions that are otherwise difficult to investigate (J. J. Russell et al., 2017).

The development of turtle organoids presented here, the first description for the chelonian Order and second for non-avian reptiles after snakes (Post et al., 2020), represents a major step towards building a toolkit that overcomes the technical challenges associated with the slow maturation and seasonality of turtle reproduction. Turtle organoids can be used to study a variety of biological processes at the cellular and molecular level, particularly as functional genomic tools become available. Namely, these turtle organoids are amenable to the implementation of techniques such as gene editing using the Clustered Regularly Interspaced Short Palindromic Repeats (CRISPR) system (Jinek et al., 2012; WareJoncas et al., 2018), providing a promising alternative for long-lived turtles over the approach used for the first genome editing of a reptile, the brown anole, *Anolis sagrei* (Rasys et al., 2019). Although successful, these editing attempts had a low throughput and required microinjections into immature oocytes of live adult females (Rasys et al., 2019), which makes large-scale editing and screening experiments impractical for turtles. The development of organoids and functional genomic tools in other turtles and reptiles overcome these limitations and will render genetic editing faster, more accessible, and more precise, and will be supported by the exponentially growing number of vertebrate genome assemblies (Rhie et al., 2021) and protocols developed here and elsewhere (Lozito et al., 2021; Post et al., 2020; Rasys et al., 2019).

### Optimization of isolation and expansion protocols for 3D turtle organoid culture

Our optimization of the organoid isolation procedures from fresh turtle liver tissue progressed first using the spiny softshell turtle (*Apalone spinifera*), then snapping turtle (*Chelydra serpentina*), and finally, hatchling, adult, and embryonic painted turtles (*Chrysemys picta*), in that order. These turtles are abundant and relatively well-studied, representing emerging models for ecology, evolution, and biomedical research (Bista et al., 2021; Das et al., 2020; Valenzuela, 2009). Turtle hepatic organoids, which were characterized and compared to the tissue of origin at the transcriptional level here, bolster the promise of these emerging species by providing a novel *in vitro* system that adds to mouse (Huch et al., 2013), human (Huch et al., 2015), and canine hepatic organoids which were successfully cultured to model human diseases (Nantasanti et al., 2015).

Some important modifications to our previously described canine organoid isolation protocol (Gabriel et al., 2022) are worth noting. The five 1X Complete Chelating Solution (CCS) washes and ethylenediaminetetraacetic acid (EDTA) incubation to separate cells from intestinal tissue and to discard the microvilli (Gabriel et al., 2022), were not required for turtle organoids, as inclusion of the tissue aided in initial growth. Therefore, our approach represents the minimal effective protocol to successfully grow turtle organoids, as described in the methods. Overall, our optimized protocol allows for expedited isolations of more samples at a lower cost while minimizing potential contamination. Incubation temperature for organoids must also follow species-specific requirements, which for the turtles studied here was 30°C, as has previously been described for 2D turtle fibroblast cultures of the same taxa (Montiel et al., 2016), whereas mammalian organoids are typically cultured at 37°C, and snake venom gland organoids at 32°C (Post et al., 2020). As previously discussed, *A. spinifera* hepatic organoids were difficult to expand in culture which led to a lower passage number for these organoid cell lines than for the other species. Hence further refinement of the protocol, media, or incubation temperature is needed to improve the yield of hepatic organoids for this species.

Because many turtle organoids exhibited holes/vacuoles in their structure which might decrease the total cell number per organoid, to increase the successful passaging and recovery rate for turtles, turtle organoids were allowed to grow larger than previously reported for canine hepatic organoids prior to passaging. Of note, some turtle liver tissues displayed dark pigmentation of the cells, which resemble melanomacrophages. Initially, these melanocytes (Schweigger & Simone, 2005) were thought to be caused by contaminants (such as bacteria and fungi) in the turtle liver organoid cultures in our study. However, the pigmented cells disappeared as leftover initial tissue was removed from the culture. Lastly, adding Prostaglandin E2 (PGE2) to the media for the culture of *C. serpentina* (*n*=2), as done for snake venom gland organoids (Post et al., 2020), allowed for successful cultivation of *C. serpentina* hepatic organoids and was therefore added to all subsequent cultures of turtle hepatic organoids.

### Characterization of turtle hepatic organoids

As expected, tissues tended to have more unique genes expressed compared with organoids, likely reflecting the greater complexity of cell types present in tissues compared with organoids. These turtle hepatic organoids are epithelial in origin and currently lack cell types such as immune cells, but emerging methods exist to co-culture immune cells with organoids (Chakrabarti et al., 2021) that may be used to bypass this limitation by creating a more complex model. Like in our turtles, *Muc5b* was also the most extreme DEG between canine hepatic organoids and canine liver tissues, as our group described elsewhere (Zdyrski et al., to be submitted), which is likely due to the fact that our media composition seems to encourage mucin production in hepatic organoids. Cholangiocytes have been shown to produce apical mucins in pig tissues and organoids (Zarei et al., 2021). Supporting the RNA expression of a mucin marker, the histology and TEM characterization of organoids identified a subset of cells in the *C. picta* hatchling organoids that secreted mucin.

Further characterization of our embryonic *C. picta* hepatic organoid model included snRNA-seq, which allows for analysis on a per cell level of resolution that is lost with bulk RNA-seq. When compared to published single-nucleus and single-cell RNA-seq data of human liver tissue (T. S. Andrews et al., 2021), our snRNA-seq revealed expression of mature cholangiocyte markers and lower expression in some clusters of some progenitor-associated cell markers. Further, our findings agree with snRNA-seq expression in human hepatic organoids of a strong cholangiocyte signature, additionally they had up upregulation of a mucin gene (MUC13) in 3D compared to 2D cultures (Aktas et al., 2022). In future experiments, specific growth factors could be used to enhance the differentiation of the culture towards a hepatocyte lineage (Huch et al., 2013), thus expanding the applicability of these hepatic organoids.

### Turtle organoids for alternative applications, such as sexual development

Our first success culturing turtle hepatic organoids opens the door to apply this technology to derive turtle organoids from other tissues and species as done by our group and others [e.g., (Bartlett et al., 2022; Post et al., 2020)], which is essential for evolutionary developmental biology and other comparative studies. For example, this emerging technology holds promise to illuminate the molecular basis of temperature-dependent sex determination (TSD), which is urgent in the face of climate change, as TSD turtle embryos develop into males or females based on ambient temperature. Indeed, the rapid increase of average temperatures on Earth threatens critical biological processes and global biodiversity (Sage, 2020). Among many detrimental effects, warmer temperatures can alter sex ratios and these skewed ratios have already threatened the viability of natural populations of several sea turtles (all TSD) (Jensen et al., 2018; Laloë et al., 2017), and a recent model predicts imminent feminization of some sea turtle populations (Hawkes et al., 2007). Yet, the full molecular basis of TSD remains obscure, despite the recent discovery of genetic and epigenetic candidates [reviewed in (Merchant-Larios et al., 2021)]. Our protocol to culture turtle organoids from *C. serpentina* (TSD) and three life stages of *C. picta* (TSD), as well as *A. spinifera,* a turtle with a ZZ/ZW sex chromosome system of genotypic sex determination (GSD) (Badenhorst et al., 2013), should be applicable, with some modification, to culture novel turtle organoids, such as gonads, to decipher the mechanisms and pathways underlying turtle sex determination.

### Benefits of 3D organoids for conservation, ecotoxicology, and evolution

Over half of all known turtle species (>50%) of data-sufficient taxa are threatened (Rhodin et al., 2018), which limits and precludes the use of these live animals for biomedical research. This limitation can be overcome by leveraging the organoid technology, a scientific toolkit that adheres to the 3Rs principles (W. M. S. Russell & Burch, 1959), while minimizing the need for tissue sampling, and thus, helps overall species survival. Carrying out research on endangered species can be challenging due to obstacles such as permit requirements, small population sizes, and low reproducibility of results, among others (Shaw et al., 2021). Being able to create a reusable *in vitro* model is therefore critical because species protection laws that protect endangered species also hinder the study of the molecular basis of adaptations of those same species, that could be essential to conservation efforts by limiting the creation of canonical transgenic models in long-lived reptiles. The *in vitro* culture of 3D organoid models from these species could offer a long-term solution to many of these inherent problems.

Turtle species can be useful biomedical models by investigating their unique adaptive strategies to overwinter, such as supercooling, anoxia tolerance, as well as their extended life spans (Bronikowski et al., 2022). 3D organoids provide a novel line of research into these remarkable adaptations, and the characterization of these organoid lines is needed prior to experimentation. Here, we utilized stranded mRNA expression analysis which revealed key differences between hatchling and adult *C. picta* hepatic organoids, including genes related to iron-binding proteins, antioxidant proteins, and serpins, which are upregulated in the liver of hatchling painted turtle in response to freezing and anoxic conditions (Storey, 2006). These organoids can accelerate the potential of turtles as a relevant biomedical model to improve human survival after stroke or heart attack, liver organ preservation before transplantation, and tissue preservation during hypoxia and anoxia.

Future studies can use this *in vitro* turtle organoid model to assess the viability of each life stage of turtles when exposed to different stressors, such as environmental toxins or xenobiotics, by using LIVE/DEAD viability and cytotoxicity staining (Forsythe et al., 2018; Votanopoulos et al., 2020). Additionally, ecotoxicological studies can be strengthened by studying organoids of sentinel turtle species that are sensitive to environmental pollutants found in their habitats. For example, sea turtle cells were used to identify potential novel biomarkers of chemical exposure, including annexin (Chaousis et al., 2021). Turtle-derived organoids could be a relevant biological model to identify such biomarkers in vulnerable populations, as they better represent the tissue of origin and harbor multiple cell types compared with conventional primary cell lines. One such important environmental pollutant in freshwater ecosystems is cadmium which can be ingested by turtles. Cadmium affects turtles’ hepatic enzyme levels, gene expression, and DNA methylation, thereby exhibiting toxic damage to the liver (Huo et al., 2020a, 2020b; Mizoguchi et al., 2022). Therefore, turtle-derived hepatic organoids could be a valuable model for studying aquatic ecosystem health and further cellular and molecular effects of heavy metal pollution on endangered and non-vulnerable species alike.

## CONCLUSION

The creation and characterization of turtle hepatic organoid lines derived from three different species of turtles, opens the door for genetic manipulation within a major vertebrate clade that is understudied due to their slow maturation, seasonality, lack of functional genomic resources, and prevalent endangered status. Turtle organoids have multiple potential applications in ecologic toxicology, supercooling/cryopreservation, and anoxia/hypoxia research, as well as many other ecological and evolutionary studies. Our study expands the application of organoid technology across the Tree of Life, facilitating future study of adaptations in reptiles and other non-model species relevant to the biological and biomedical communities.

## METHODS

### Animal husbandry and tissue collection

Five juvenile spiny softshell turtles (*Apalone spinifera*), three juvenile snapping turtles (*Chelydra serpentina*), plus two embryonic, three hatchlings, and three adult painted turtles (*Chrysemys picta*) were used in this study. Animals were collected in Iowa under appropriate permits from the Iowa DNR (SC648 and SC595), and all procedures followed protocols approved by the Institutional Animal Care and Use Committee (IACUC) (IACUC-21-121) of Iowa State University as described below. Details of the donor animals, including sex, age, and the outcome of the organoid culture, can be found in **Table S1**. Adult males and freshly laid *C. picta* eggs were collected from the wild. Hatchlings and juveniles were obtained from eggs incubated in the laboratory at 26°C (*C. picta*), a temperature that produces exclusively males in painted turtles, as this species displays temperature-dependent sex determination (TSD). The embryonic sample at developmental stage 22 (Yntema, 1968), was obtained from a *C. picta* egg incubated at 26°C. Eggs of *C. serpentina* (TSD) were incubated at 27.5°C, which produces a mixed sex ratio (Ewert et al., 1994), such that sex of snapping turtle juveniles was diagnosed by gonadal inspection. In contrast, *A. spinifera* displays a ZZ/ZW sex chromosome system of genotypic sex determination (GSD) (Badenhorst et al., 2013) and produces both sexes at 27.5°C (Bull & Vogt, 1979), the temperature used here to incubate their eggs. Thus, *A. spinifera* juveniles can be sexed by PCR amplification of sex-linked markers (Literman et al., 2017), a simpler method than by qPCR of rDNA repeats (Literman et al., 2014).

Live animals were housed indoors in water tubs, provided with UV A/B bulbs and a dry surface for basking, and kept at ∼24°C until processing. Animals were fed Tetra ReptoMin sticks *ad libitum*. Animals were euthanized, then washed in iodine and hydrogen peroxide, and sex was diagnosed (*C. serpentina*) or confirmed (*C. picta* and *A. spinifera*) by gross gonadal morphology or presumed by the incubation temperature (*C. picta* embryo). Tissues were quickly harvested inside a biosafety cabinet, and a subset of tissue was immediately placed into RNAlater (Invitrogen; AM7021) (except embryonic liver) and another in formalin for downstream processing.

### Organoid culture

Experimental methods for liver organoid isolation followed our previously published canine protocol (Gabriel et al., 2022) with minor modifications. The formulation of expansion media (*Complete media with growth factors with ROCK inhibitor and GSK3*β *inhibitor - CMGF+ R/G*) and the optimization and variations of the organoid isolation protocol are listed in **Tables S3 and S1** respectively. The optimized and minimum required protocol for organoid hepatic isolation consisted of rinsing the fresh tissue in Phosphate Buffered Saline (PBS)/N-acetylcysteine (NAC) once and then transferring it to a tube filled with Advanced DMEM/F12 (Gibco; 12634-010), mincing the tissue into small fragments, washing in complete chelating solution (1X CCS) once, vortexing of the sample, removing the supernatant, adding 6 mL of DMEM, centrifuging at 100 g (700 g was used for the first samples, but was lowered to 100 g to assist in the separation of dead cells or debris) for 5 min at 4°C, removing the supernatant, mixing the pellet of cells and tissue fragments (to aid in initial growth) with Matrigel® Matrix (Corning; 356231, 356255), subsequent plating in 24 well plates (Corning; 3524) and adding media (**Fig. 1*A***). Because embryonic *C. picta* liver tissue was smaller than at later life stages, the protocol was further shortened to submersion in DMEM, centrifuging at 100 g for 5 min at 4°C, removal of supernatant, addition of 200 µL DMEM, centrifuging again, mincing, transfering to a tube and centrifuging, then mixing with Matrigel and plating. Prostaglandin E2 (PGE2) (Tocris; 2296) 9.93 nM was added to the culture media, as was previously done for snake venom gland organoids (Post et al., 2020). When passaging or cleaning, organoids were resuspended in Matrigel solidified at 37°C for ∼15-30 minutes to avoid organoid damage at warmer temperatures. The passaging technique used consisted of adding 500 µL of TrypLE™ Express (Gibco; 12604-021) to ∼500 µL of organoid resuspended in DMEM at 37°C for 10 min, with gentle flicking halfway through. When culturing embryonic hepatic organoids and re-growing frozen samples, if organoids would not pellet, the addition of 2 mL of Cell Recovery Solution (Corning; 354270) to 1 mL of organoids suspended in DMEM and a subsequent incubation on ice for 10 minutes assisted in degrading excess Matrigel. After spinning, 6 mL of DMEM was used to wash away the Cell Recovery Solution before plating. Additionally, if samples previously resisted passaging when given TrypLE™ Express, samples were passed in Cell Recovery Solution as opposed to DMEM which typically dissociated the organoids into single cells or small clusters. Organoids were incubated in a Nuaire Direct Thermal incubator at 30°C and 5% CO_2_.

### Biobank creation

Organoids were frozen in alternative freezing media, including (1) 50% CMGF+ R/G + PGE2, 40% FBS, and 10% DMSO as well as (2) Cryostor CS10 (Stem Cell; 210102) to test their biobanking potential. Before freezing, organoids were resuspended in freezing media, placed overnight at −80°C in a Mr. Frosty container (Thermo Fisher Scientific; 5100-0001) filled with isopropanol, and then transferred to liquid nitrogen (−196°C) indefinitely.

### Transmission electron microscopy

Samples were preserved in 1% paraformaldehyde and 3% glutaraldehyde in phosphate-buffered saline (PBS) and stored at 4°C prior to processing. The samples were fixed for 48 hours at 4°C in 1% paraformaldehyde, 3% glutaraldehyde in 0.1 M sodium cacodylate buffer, pH 7.2. After washing in a cacodylate buffer 3 times for 10 minutes each, samples were post-fixed with 1% osmium tetroxide in 0.1M sodium cacodylate buffer at room temperature for 1 hour. Next, the samples were washed with deionized water 3 times for 15 minutes each, and then *en bloc* stained using 2% uranyl acetate in distilled water for 1 hour. Then the samples were washed for 10 minutes in distilled water, dehydrated for 1 hour in each step of a graded ethanol series (25, 50, 70, 85, 95, 100%), dehydrated further with 3 changes of pure acetone for 15 minutes each, and then infiltrated with EmBed 812 formula (hard) for EPON epoxy resin (Electron Microscopy Sciences, Hatfield, PA) with graded ratios of resin to acetone until fully infiltrated with pure epoxy resin (3:1, 1:1, 1:3, pure) for 6-12 hours per step. The tissues were then placed into BEEM capsules and polymerized at 70°C for 48 hours. Then, using a Leica UC6 ultramicrotome (Leica Microsystems, Buffalo Grove, IL), 1.5 µm thick sections were made and stained with EMS Epoxy stain (a blend of toluidine blue-O and basic fuchsin). Thin sections were made at 50 nm and collected onto single slot carbon film grids. Finally, TEM images were captured using a 200 kV JEOL JSM 2100 scanning transmission electron microscope (Japan Electron Optics Laboratories, Peabody, MA) with a GATAN One View 4K camera (Gatan inc., Pleasanton, CA).

### Histological stains

Tissue samples were fixed in 10% formalin while organoids had their media removed and 500 µL of Formalin-acetic acid-alcohol (FAA, composition in Gabriel et al. 2022) was added to wells containing Matrigel (Gabriel et al., 2022). Both tissues and organoids were changed to 70% ethanol 24 hours later prior to paraffin-embedding. Tissue and organoids were stained with hematoxylin and eosin (H&E), Alcian Blue, and Periodic acid–Schiff (PAS) at the Iowa State University Histopathology Department. After staining, slides were scanned on Leica Aperio GT 450 Scanner and analyzed with ImageScope (v12.4.3.5008) to characterize the morphology and cell type composition of tissues and organoids.

### RNA extractions

Expanded organoids were transferred to a 15 mL tube with Advanced DMEM/F12, centrifuged at 700 g at 4°C for 5 min, resuspended in Cell Recovery Solution, placed in ice for ∼10 min to degrade Matrigel, centrifuged, the supernatant was discarded, and the pellet was resuspended in 100 µL of PBS and transferred to a cryovial. A volume of 900 µL of RNAlater was used to flush the 15 mL tube and then added to the cryovial before storage in either liquid nitrogen (−196°C) or a −80°C freezer. Tissue biopsies were stored in liquid nitrogen or at −80°C in cryovials containing 1 mL of RNAlater. After thawing, tissues were rinsed in PBS to remove excess RNAlater solution, immediately transferred into 800 µL of Trizol (Life Technologies; 15596026) and homogenized with a pestle. Organoids were thawed, transferred to a 15 mL tube containing 2 mL of PBS (Corning; 21-040-CM) and centrifuged at 1,200 g at 4°C for 5 min to pellet. Excess RNAlater was removed, 1 mL of Trizol was added to the organoids, and then briefly vortexed (5-10 seconds).

Homogenized samples were stored at room temperature for 5 minutes, then centrifuged for 10 minutes at 12,000 g at 4°C to eliminate debris and polysaccharides, whereas the supernatant was transferred to a new tube. Chloroform (Alfa Aesar; J67241) was added (0.2 mL chloroform per mL Trizol), and samples were vigorously shaken for 20 seconds and incubated at room temperature for 2-3 minutes. Samples were centrifuged at 10,000 g for 18 minutes at 4°C, and the aqueous phase was transferred into a new sterile 1.5 mL RNase-free tube. Then, an equal volume of 100% RNA-free EtOH was slowly added using a pipette, mixed, then the samples were transferred to a Qiagen RNeasy column (RNeasy Mini kit; Qiagen; 74104) seated in a collection tube, and centrifuged for 30 seconds at 8,000 g, after which the flow-through was discarded and the Qiagen DNase treatment protocol was followed. Next, 500 µL of buffer RPE was added and samples were centrifuged for 30 seconds at 8,000 g. After discarding the flow-through, another 500 µL of buffer RPE was added and samples were centrifuged for 2 minutes at 8,000 g. Flow-through was discarded, and columns were centrifuged for 1 minute at 8,000 g to remove the remaining buffer. RNA was eluted in 50 µL of RNase-free water (Sigma; W4502-50ML) and allowed to sit for 2 minutes before centrifuging for 1 minute at 8,000 g. Samples were centrifuged again at 8,000 g, immediately analyzed on a Nanodrop ND-1000 Spectrophotometer (Thermo Fisher Scientific), and stored at −80°C.

### RNA sequencing

RNA samples were shipped to GENEWIZ for analysis as follows. The Qubit 2.0 Fluorometer (ThermoFisher Scientific, Waltham, MA, USA) was used to quantify RNA concentration, and a 4200 TapeStation (Agilent Technologies, Palo Alto, CA, USA) was used to measure RNA integrity. Next, an ERCC RNA Spike-In Mix kit (ThermoFisher Scientific; 4456740) was added to normalize the total RNA prior to library preparation. A strand-specific RNA sequencing library was prepared using NEBNext Ultra II Directional RNA Library Prep Kit for Illumina (NEB, Ipswich, MA, USA). Then, the enriched RNAs were fragmented at 94°C for 8 minutes. First-strand and second-strand cDNA were subsequently synthesized, with the second strand of cDNA marked by incorporating dUTP during the synthesis (which quenched the amplification of the second strand, helping preserve the strand specificity). cDNA fragments were adenylated at 3’ ends, and an indexed adapter was ligated to cDNA fragments, with a limited cycle PCR being used for library enrichment. The sequencing library was then validated on the Agilent TapeStation and quantified using a Qubit 2.0 Fluorometer in addition to quantitative PCR (KAPA Biosystems, Wilmington, MA, USA). The sequencing libraries were multiplexed and clustered onto two flowcells, loaded onto an Illumina HiSeq 4000 instrument, according to the manufacturer’s instructions, and sequenced using a 2×150bp Paired-End (PE) configuration. The HiSeq Control Software (HCS) was used to conduct image analysis and base calling. Raw sequence data (.bcl files) generated from Illumina HiSeq was converted into fastq files and de-multiplexed using Illumina bcl2fastq 2.20 software with one mismatch allowed for index sequence identification.

### Transcriptome assembly

Reads were trimmed with trimmomatic (version 0.39) (Bolger et al., 2014) to remove low-quality bases and adapter contamination. Reads were checked post-trimming with FASTQC (v 0.11.7) (S. Andrews, 2010) to confirm quality. Following trimming, reads were mapped to the *C. picta* RefSeq genome (Chrysemys_picta_BioNano-3.0.4) (Lee et al., 2020) using GSNAP (version 2021-03-08) (Wu & Nacu, 2010; Wu & Watanabe, 2005). Read representation was calculated using samtools (version 1.10) (Li et al., 2009). Following mapping, individual library BAM files were genome-guided assembled with StringTie (version 1.3.4a) (Pertea et al., 2015) and then merged into a single assembly using the --merge function. Following merging, transcript abundances were calculated for each library and counts were extracted using the prepDE.py script.

### Spike-in and differential expression analysis

In parallel, ERCC reads were mapped to the ERCC reference following the same assembly pipeline as for the sample reads, except that discovery of novel transcripts was not permitted during assembly. Counts for the ERCC transcripts were appended to the gene count matrix for *C. picta*. Differential expression of gene models was calculated with DESeq2 (version 1.24.0) (Love et al., 2014) in R (version 3.6.2) (R Core Team, 2018), testing for the effect of age, sample type and their interaction via a full factorial ANOVA (model Y ∼ Age * Sample Type). Estimation of size factors was based on ERCC spike-in transcripts for normalization of the data. As many genes showed an interaction effect, the full factorial model was retained. Differentially expressed genes were filtered based on a baseMean (mean of the counts for all samples that have been normalized for sequencing depth) of > 50 and a *P*-adjusted value < 0.05.

### Annotation and enrichment analysis

Blastx (blast-plus version 2.7.1) (Camacho et al., 2009) against the Uniprot database (accessed May 24, 2022) (Consortium, 2019) was used to further annotate transcripts that were not annotated during the initial genome-guided assembly, although some transcripts remained unannotated after this blastx. Transcript sequences were extracted from the transcriptome using gffread (v0.12.7) (Pertea & Pertea, 2020). These transcripts were then translated into peptide sequences using TransDecoder (version 5.5.0; https://github.com/TransDecoder/TransDecoder). These sequences were then searched against the PANTHERDB (v17.0) (Mi et al., 2019; Thomas et al., 2022) hidden Markov models to obtain compatible sequence identifiers for enrichment analysis. Stringtie transcript counts were converted to gene-level lengthScaledTPM using the tximport package (v1.18.1) (Soneson et al., 2015) in R. Genes were then mapped to their corresponding PANTHER IDs via transcript isoforms. In the case where multiple isoforms were present, the isoform with the best supported PANTHER ID was prioritized. Unannotated transcripts were filtered out of the analysis, as this was required by the program.

PANTHER IDs and corresponding expression values were submitted to pantherdb.org for statistical enrichment analysis (Released 2022-10-17) which uses a Mann-Whitney U test to calculate enrichment of GO terms. Enrichment analysis was performed for each library and was searched against the following databases (v17.0 Released 2022-02-22) (Ashburner et al., 2000; Carbon et al., 2021): Pathways, GO-Slim Molecular Function, GO-Slim Biological Process, GO-Slim Cellular Component, and Protein Class. GO terms were filtered for terms that were over-enriched (as opposed to under-enriched). Following filtering, terms were filtered for those present in all three biological replicates. These over-enriched and replicated terms were input into REVIGO (Supek et al., 2011) for visualization. REVIGO reduces redundancy in lists of GO terms by considering semantic similarity and identifies terms that are most representative of clusters of related terms. It then allows the user to generate graphical representations of these relationships to aid interpretation of the enrichment results. REVIGO was run with a cutoff of 0.5 and terms were provided with FDR values. Obsolete terms were removed, and the dataset was compared to the Whole Uniprot Database and used the SimRel semantic similarity measure. Analyses were run on 2022/12/09 and the databases used for reference were go.obo (2022-11-03) and goa_uniprot_gcrp.gaf.gz (2022-09-16). Treemap (Tennekes & Ellis, 2017) was used to visualize the resulting clusters. The REVIGO-provided Rscript was downloaded and used to generate plots for interpretation.

### Single-nuclei RNA-seq harvesting

For single-nuclei processing, organoids were harvested from a sample that had undergone multiple passages but was not previously frozen. Media was removed and 500 µL of Cell Recovery Solution was added to each well, mixed, then placed in ice for 20 minutes to degrade Matrigel. The sample was centrifuged at 100 g for 5 minutes at 4°C, supernatant was removed, and the pellet was washed with approximately 6 mL of DMEM and centrifuged at 100 g at 4°C for 5 min. The organoid pellet was resuspended in 500 µL of DMEM and transferred to a cryovial. The cryovial was centrifuged at 100 g at 4°C for 5 min, all supernatant was removed, and the pellet was flash frozen in liquid nitrogen for approximately 30 seconds. The sample was then immediately stored at −80°C, shipped on dry ice the following day to Azenta (South Plainfield, NJ, USA), and stored at −80°C prior to processing as described below.

### Nuclei isolation

Nuclei extraction was performed using the Miltenyi Nuclei Extraction Buffer (Miltenyi Biotec, Auburn, CA, USA) following manufacturer’s guidelines with gentle MACS Dissociation and C tubes. Upon isolation, the nuclei were counted using trypan blue with a Thermo Fisher Countess III automated cell counter.

### 3’ RNA library preparation and sequencing

Single nuclei RNA libraries were generated using the Chromium Single Cell 3’ kit (10X Genomics, CA, USA). Loading onto the Chromium Controller was performed to target capture of ∼10,000 GEMs per sample for downstream analysis and processed through the Chromium Controller. Quality of the sequencing libraries were evaluated on the Agilent TapeStation, then quantified using a Qubit 2.0 Fluorometer (Invitrogen, Carlsbad, CA). Prior to loading onto an Illumina sequencing platform, pooled libraries were quantified using qPCR (Applied Biosystems, Carlsbad, CA, USA). The samples were sequenced at a configuration compatible with the recommended guidelines outlined by 10X Genomics. Raw sequence data (.bcl files) were converted into fastq files and de-multiplexed using the 10X Genomics’ cellranger mkfastq command. Subsequent UMI and cell barcode de-convolution along with mapping to the reference genome Chrysemys_picta_bellii-3.0.3 (GCA_000241765.2) were performed using 10X Genomics Cell Ranger 6.0.1 (Zheng et al., 2017) software package to generate the final digital gene expression matrices and cloupe files.

### Analysis of single-nuclei RNA sequencing data

snRNA-seq reads were mapped to the reference genome (Chrysemys_picta_BioNano-3.0.4) then the Seurat package (v. 4.0) (Hao et al., 2021) in R (v. 4.2) (R Core Team, 2018) was used. The percentage of reads mapping to mitochondrial genes was used to minimize mitochondrial contamination typically seen in low quality or dying cells, and those with a high percentage were filtered out. Cells were filtered to retain those with gene counts between 200 and 6000 and having less than 40% mitochondrial contamination. Next, the data was normalized using the log Normalization method. Then features with high cell to cell variation were identified. The data was then scaled so that the mean expressions across the cells were 0 and variance across the cells was 1 prior to performing a principal component analysis. Clustering of samples utilized the KNN method, and using default parameters, clusters were identified.

## ACKNOWLEDGMENTS

We thank Dr. Amanda Kreuder for the use of her incubator during this experiment. We thank Irah Dhaseleer (https://dhaseleerillustration.com/) and Regan Schmidt (https://regan264.wixsite.com/regan-schmidt) for their illustrations. We wish to thank Dr. Olufemi Fasina and Dr. Austin Viall for their expertise in identifying pathological features. We thank Basant Ahmed for her help with the project. We thank David Fernández and Vanessa Livania for their help in sample maintenance and documentation. We appreciate the timely processing of samples at the Iowa State University Histology department and the Iowa State University Pathology department. We wish to thank Tracey Stewart for processing and imaging TEM samples.

## DATA AVAILABILITY

The RNA-seq stranded mRNA raw reads and snRNA-seq reads generated and analyzed in this study are available in the Sequence Read Archive (NCBI-SRA BioProject PRJNA931617). Bioinformatic scripts are available on Github (https://github.com/chris-zdyrski/Turtle_Hepatic_Organoids).

## SUPPLEMENTAL

**Biological Processes of turtle liver and hepatic organoids.**

**Supplemental Fig. 1:** The Biological Processes identified when analyzing the transcriptomic expression patterns of liver tissue and hepatic organoids. Enrichment analysis of (A) hatchling tissue (*n*=3), (B) adult tissue (*n*=3) samples of liver in addition to hepatic cultures of (C) hatchling organoid (*n*=3), (D) adult organoid (*n*=3) of *C. picta* hepatic samples.

**Cellular Components of turtle liver and hepatic organoids.**

**Supplemental Fig. 2:** The Cellular Components identified when analyzing the transcriptomic expression patterns of liver tissue and hepatic organoids. Enrichment analysis of (A) hatchling tissue (*n*=3), (B) adult tissue (*n*=3) samples of liver in addition to hepatic cultures of (C) hatchling organoid (*n*=3), (D) adult organoid (*n*=3) of *C. picta* hepatic samples.

**Molecular Functions of turtle liver and hepatic organoids.**

**Supplemental Fig. 3:** The Molecular Functions identified when analyzing the transcriptomic expression patterns of liver tissue and hepatic organoids. Enrichment analysis of (A) hatchling tissue (*n*=3), (B) adult tissue (*n*=3) samples of liver in addition to hepatic cultures of (C) hatchling organoid (*n*=3), (D) adult organoid (*n*=3) of *C. picta* hepatic samples.

**Histopathology staining of softshell turtle hepatic tissues.**

**Supplemental Fig. 4:** Overview of turtle liver histology from the spiny softshell turtle (*Apalone spinifera*). (A-C) Key histologic landmarks identified using hematoxylin and eosin (H&E) in the turtle liver tissue include the portal venule (Pv) and terminal hepatic venule (Hv). In the portal region, a small bile duct is seen (B, inset and arrow). (D) Note the large intrahepatic bile duct (inset). (E) Nucleated erythrocytes are seen within sinusoids (red arrows) and melanomacrophages (black arrow) are readily detected by their cytoplasmic pigment (black arrows). Hepatocytes often have round nuclei that are eccentrically and basolaterally located in the cytoplasm near the sinusoidal border. Hepatocytes (black arrowheads) are distended and have increased rarefaction of the cytoplasm, which parallels the presence of PAS+ glycogen (E, inset, magenta coloration).

**Supplemental Table 1:** Sample information for attempted organoid cultures including ID, species, age, and sex. Passage number and outcomes for all attempted turtle organoid cultures. Details on which samples were harvested for characterization and inclusion into the biobank are listed.

**Supplemental Table 2:** Stranded RNA-seq reads mapped per sample for both tissue and organoids.

**Supplemental Table 3:** Media composition for CMGF+ used in the optimized cultures as well as concentrations, manufacturers, and additional growth factors are listed.

