## Supplementary figures and images for "Characterization of the First Turtle Organoids: A Model for Investigating Unique Adaptations with Biomedical Potential"

### Supplemental Figure 1

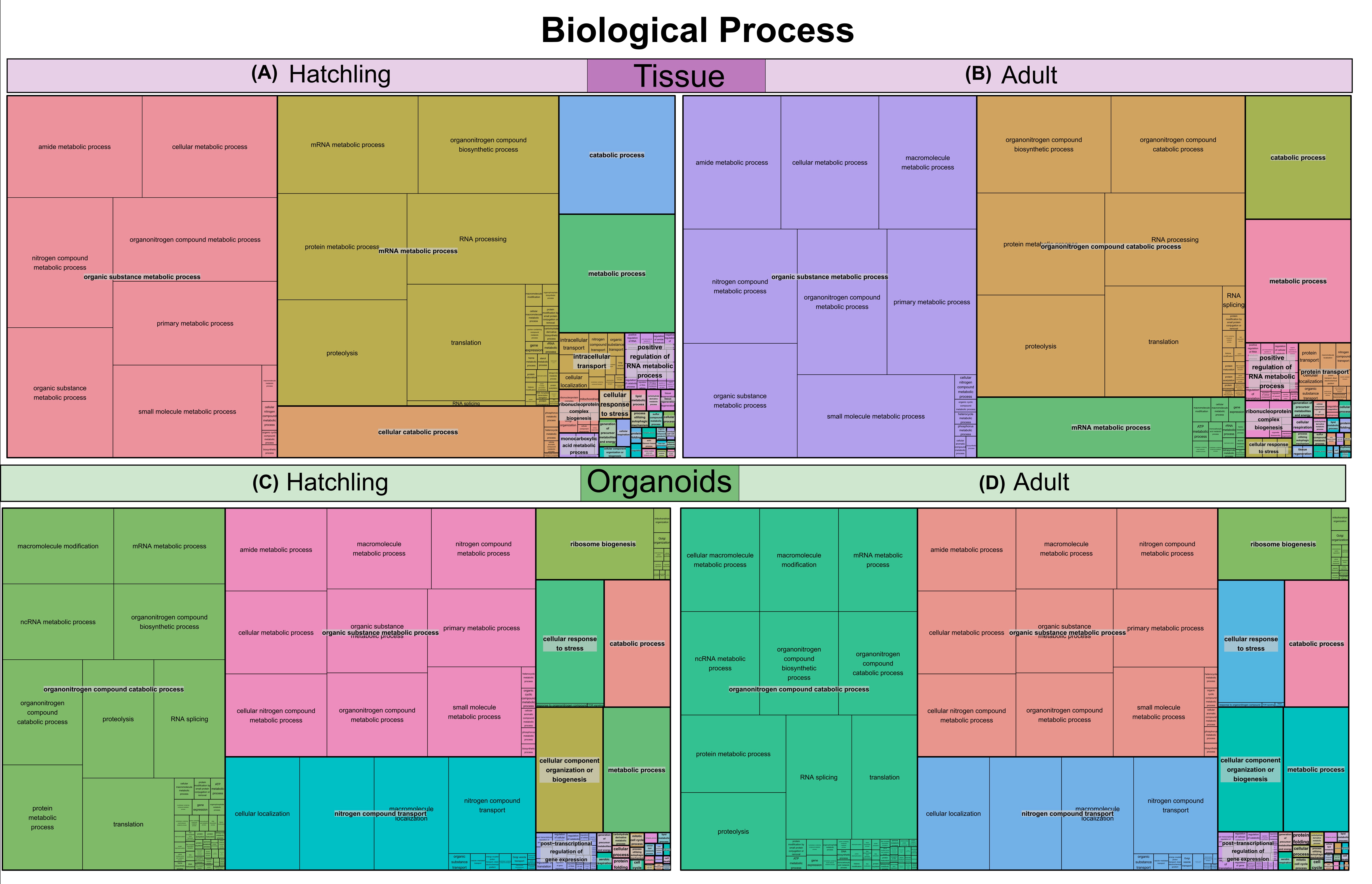

### Supplemental Figure 2

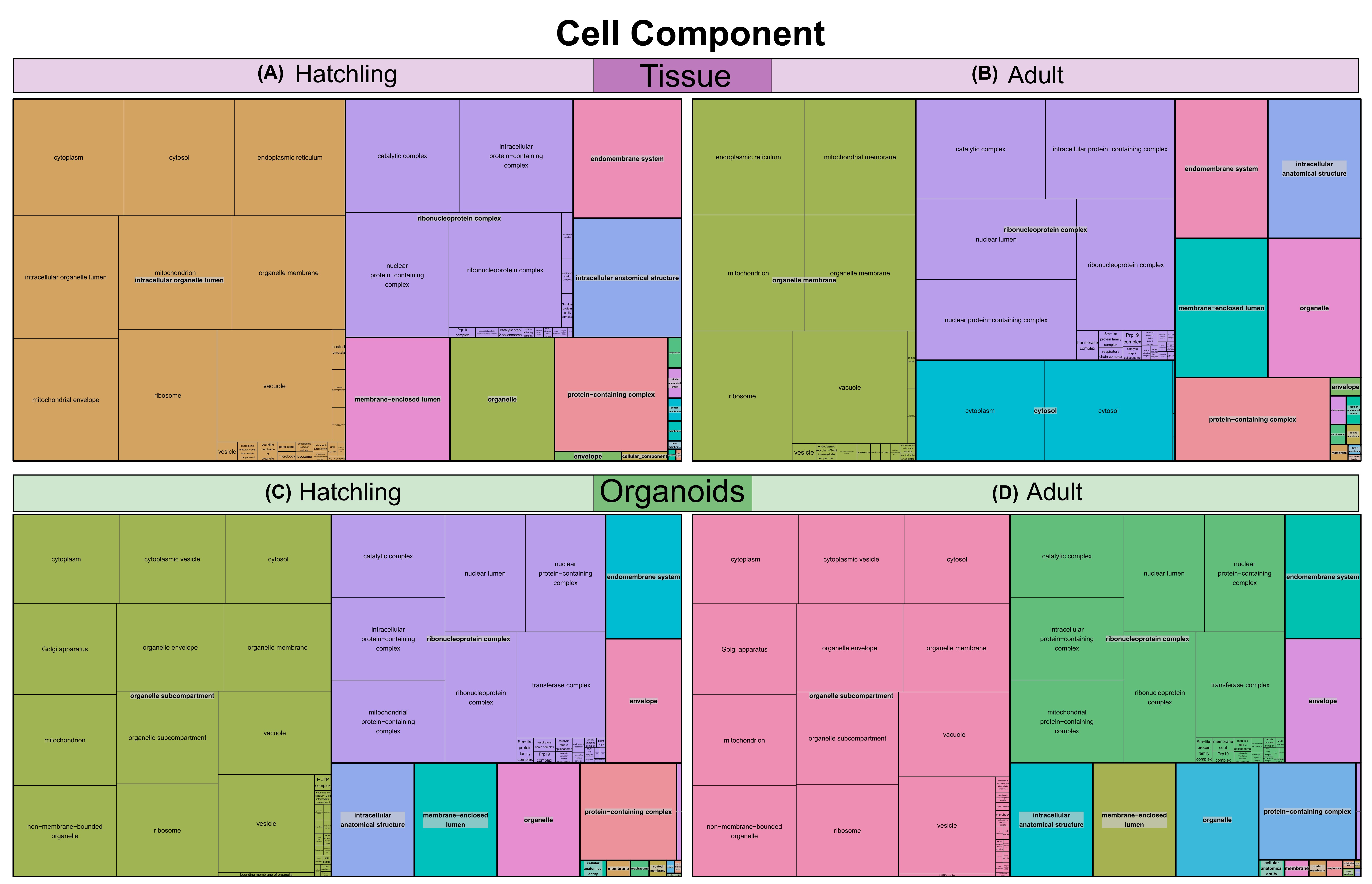

### Supplemental Figure 3

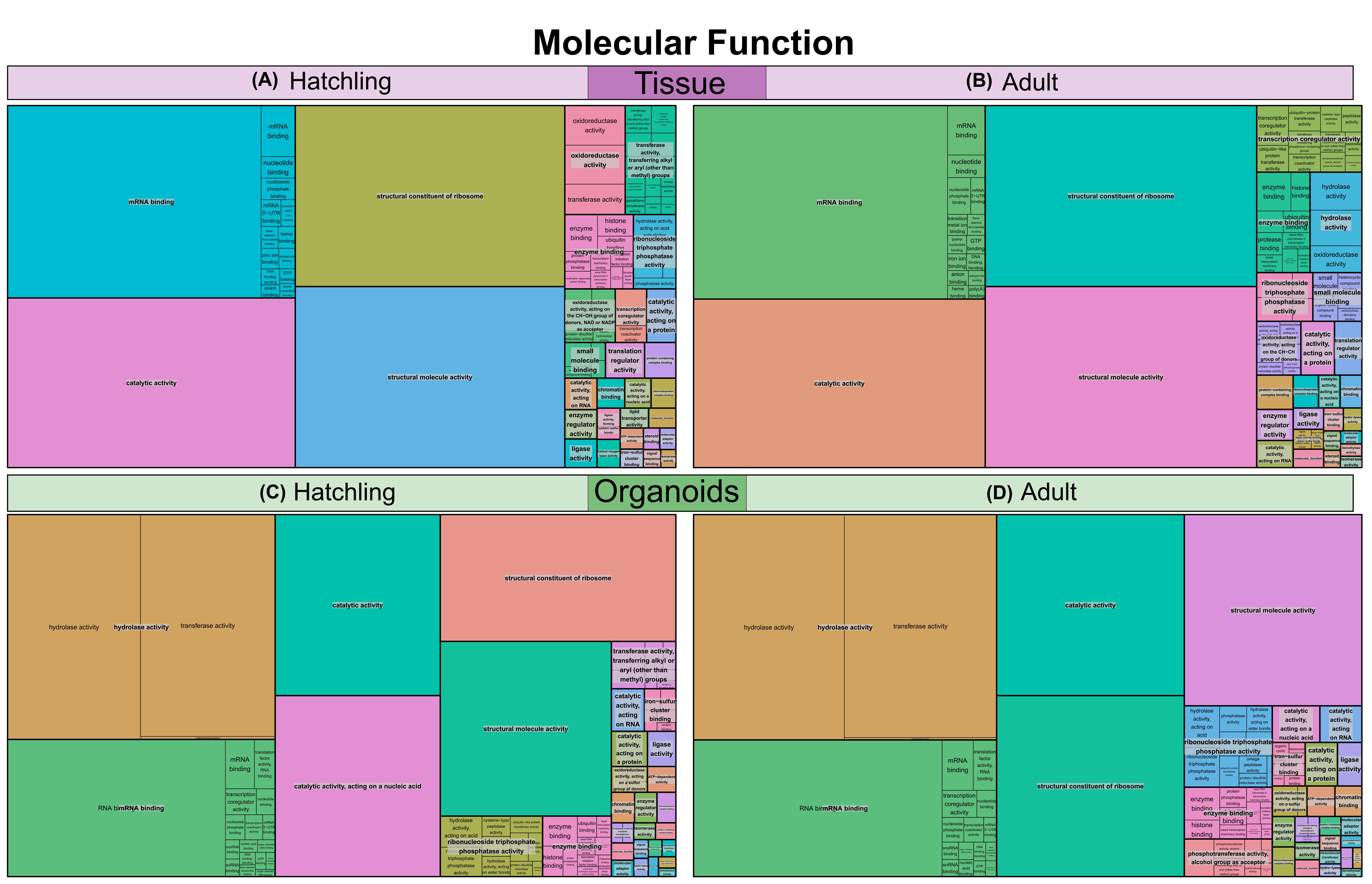

### Supplemental Figure 4

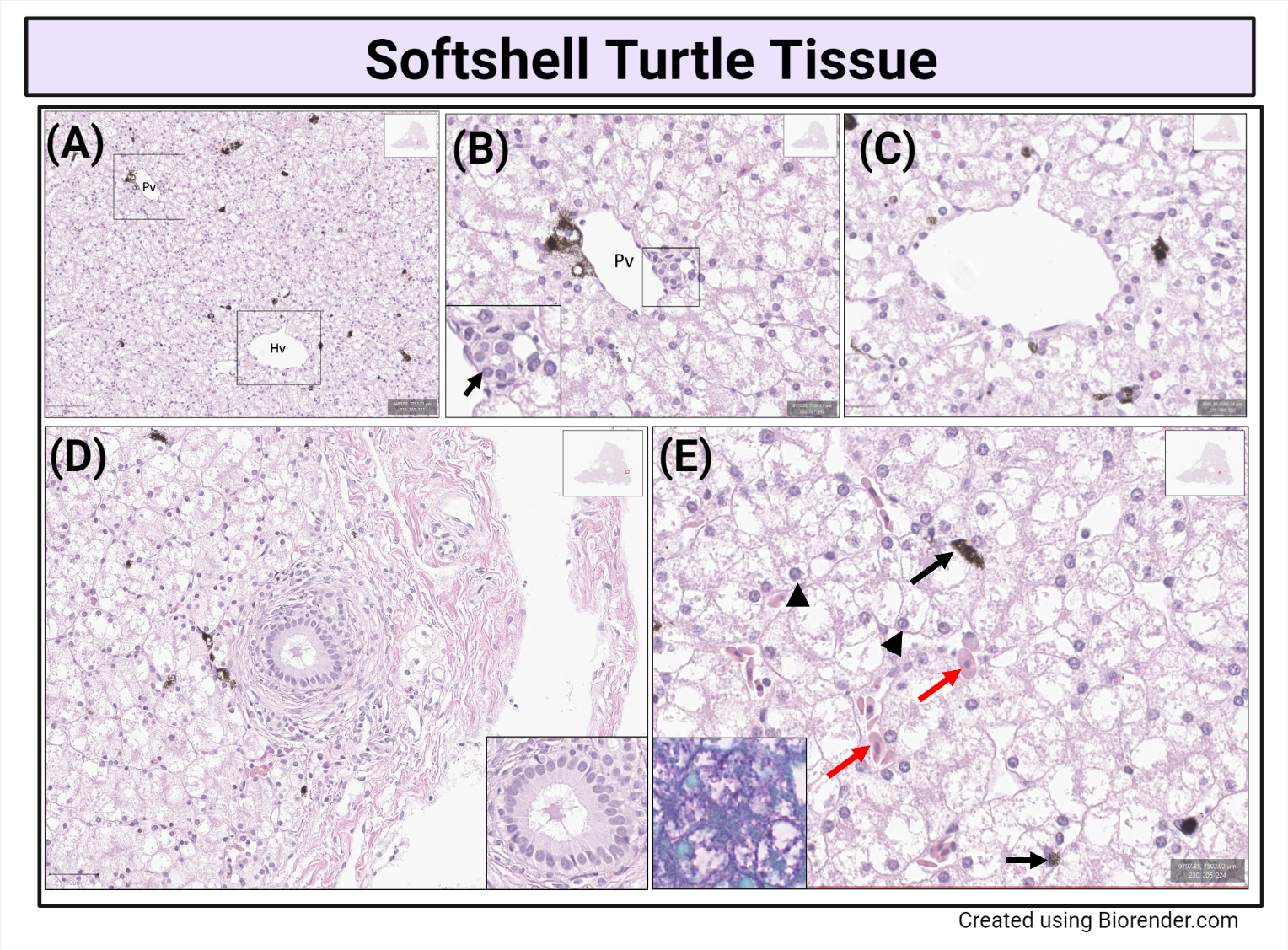
